# A nanocompartment containing the peroxidase DypB contributes to defense against oxidative stress in *M. tuberculosis*

**DOI:** 10.1101/2020.08.31.276014

**Authors:** Katie A. Lien, Robert J. Nichols, Caleb Cassidy-Amstutz, Kayla Dinshaw, Matthew Knight, Rahul Singh, Lindsay D. Eltis, David F. Savage, Sarah A. Stanley

## Abstract

Encapsulin nanocompartments are an emerging class of prokaryotic protein-based organelles consisting of an encapsulin protein shell that encloses a protein cargo^1^. Genes encoding nanocompartments are widespread in bacteria and archaea, and recent works have characterized the biochemical function of several cargo enzymes^2^. However, the importance of these organelles to host physiology is poorly understood. Here, we report that the human pathogen *Mycobacterium tuberculosis* (Mtb) produces a nanocompartment that contains the dye-decolorizing peroxidase DypB. We show that this nanocompartment is important for the ability of Mtb to resist oxidative stress in low pH environments, including during infection of host cells and upon treatment with a clinically relevant antibiotic. Our findings are the first to implicate a nanocompartment in bacterial pathogenesis and reveal a new mechanism that Mtb uses to combat oxidative stress.

## Results

Bacterial cells were long thought to lack compartmentalization of function. However, the identification of organelle-like structures including microcompartments, anammoxosomes, magnetosomes and, most recently, encapsulin nanocompartments have revolutionized our understanding of bacterial cell biology^3–5^. Characterized nanocompartments are proteinaceous shells that are 24 to 45 nm in diameter and comprised of 60-240 subunits of a single protomer^1,6^. This shell surrounds (“encapsulates”) an enzymatic cargo protein^1^. Although putative encapsulin systems have been identified in >900 bacterial and archaeal genomes^2,7^, very little is known about their physiological function. Based on genomic organization, encapsulin systems are often predicted to compartmentalize enzymes involved in oxidative stress defense, iron storage, and anaerobic ammonium oxidation^2^. However, the function of an encapsulin system has only ever been demonstrated in a single bacterial species, *Myxococcus xanthus*^8^, and the physiological role of nanocompartments is largely unexplored.

The *Mycobacterium tuberculosis* genome encodes the predicted encapsulin gene *Rv0798c/*Cfp29^9^ in a two-gene operon with *Rv0799c*, the dye-decolorizing peroxidase DypB (Figure 1A). Overexpression of the predicted Mtb encapsulin gene in *Escherichia coli* was previously shown to result in the formation of nanocompartment-like structures^9^. Three potential cargo proteins for the nanocompartment were proposed based on a putative shared encapsulation targeting sequence: DypB, FolB, and BrfB. In *E. coli*, overexpression of each protein with Cfp29 resulted in encapsulation. However, this study did not address whether Mtb produces endogenous nanocompartments or identify the specific function of these compartments in Mtb biology. A transposon screen has identified Cfp29 as a gene required for growth in mice^10^ and Cfp29 has long been known as an immunodominant T cell antigen in both mice and human TB patients^11^. Taken together, these results suggest that Mtb may produce an encapsulin nanocompartment that is important for infection.

**Figure 1.**
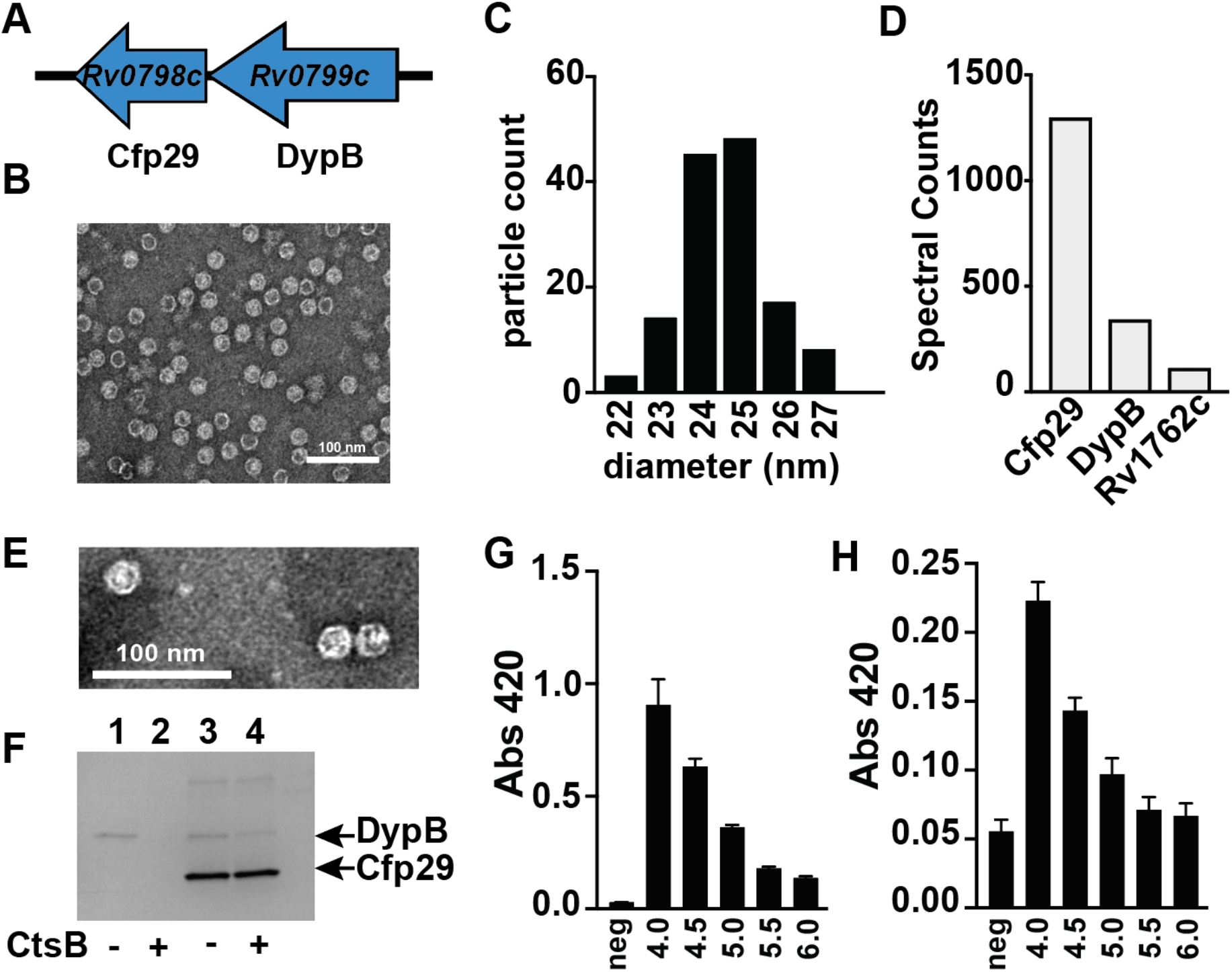
Mtb produces endogenous nanocompartments that package a peroxidase. **(A)** Schematic of the nanocompartment operon in Mtb which encodes the encapsulin shell protein (Cfp29) and the dye-decoloring peroxidase cargo protein (DypB). **(B)** TEM of Cfp29 encapsulin proteins purified following heterologous expression of the Mtb nanocompartment operon in *E. coli.* **(C)** Size distribution of Cfp29 protomers purified from *E. coli*. **(D)** Peptide counts from mass spectrometry analysis of endogenous nanocompartments purified from Mtb. **(E)** TEM of endogenous nanocompartments purified from Mtb. **(F)** Coomassie-stained SDS-PAGE of unencapsulated (lanes 1 and 2) and encapsulated (lanes 3 and 4) DypB following 14 hours of Cathepsin B (CtsB) treatment. Peroxidase activity of **(G)** unencapsulated and **(H)** encapsulated DypB (5 nM) using ABTS (480 nM) as a substrate in the presence of H_2_O_2_ (480 nM) at varying pH levels (4.0-6.0) as reported by a change in the absorbance at 420 nm. neg=no added enzyme.

To confirm the previous finding that heterologous expression of *Rv0798c* and *Rv0799c* in a host species results in the assembly of an encapsulin system, we expressed these genes in *E. coli* and isolated nanocompartments. Clarified protein lysates from *E. coli* were purified by ultracentrifugation and size exclusion chromatography. Assembled encapsulin nanocompartments are distinguishable by their high molecular weight^10^. Indeed, a fraction from the purification contained a high-molecular weight species >260 kDa observable on an SDS-PAGE gel (Figure S1A). Fractions containing putative nanocompartments were pooled and imaged using transmission electron microscopy, which revealed the presence of nanocompartment-like icosahedral structures with the expected diameter of ~25 nm (Figure 1B, 1C).

To determine whether Mtb produces nanocompartments under normal laboratory growth conditions, we performed an ultracentrifugation-based nanocompartment isolation strategy using wild-type H37Rv strain bacteria grown to mid-log phase (Figure S1B). Mass spectrometry analysis of the nanocompartment fraction identified both the encapsulin protein Cfp29 and the peroxidase DypB (Figure 1D). TEM analysis confirmed the presence of nanocompartment particles ~25 nm in diameter (Figure 1E). Interestingly, Rv1762c, a protein of unknown function, was consistently identified in purified nanocompartment preparations from Mtb (Figure 1D). We were unable to identify either FolB or BrfB in nanocompartments from Mtb, suggesting that these proteins are not endogenous substrates for encapsulation under normal laboratory growth conditions for Mtb.

Cfp29 (‘<u>c</u>ulture <u>f</u>iltrate <u>p</u>rotein 29’) was originally identified in the supernatants of Mtb cells grown in axenic culture^12^. As Cfp29 lacks a secretory signal sequence and is part of a large macromolecular complex, it is unclear how a nanocompartment could be actively secreted. However, nanocompartment structures are remarkably stable^13^ and it is possible that nanocompartments released from dying bacteria accumulate in culture as they are highly resistant to proteolysis/degradation. To test this hypothesis, we purified unencapsulated DypB and encapsulated DypB and exposed them to a lysosomal protease, cathepsin B, while monitoring proteolysis. Whereas unencapsulated DypB was completely degraded after 14 hours of exposure to cathepsin B, encapsulated DypB was protected from degradation, demonstrating the resistance of nanocompartment structures to proteolysis (Figure 1F).

DypB proteins are known to have low pH optima^9,14^. To demonstrate that Mtb DypB has a similarly low pH optimum, SUMO-tagged unencapsulated DypB was purified from *E. coli* (Figure S2A), and the peroxidase activity was evaluated at a range of pH values using the ABTS (2,2’-azino-bis(3-ethylbenzothiazoline-6-sulfonic acid)) dye-decolorizing assay previously used to characterize DypB proteins^9,15^. Similar to the *Vibrio cholerae* DypB^14^, purified Mtb DypB had increased enzymatic activity in low pH environments, with the greatest efficacy at pH 4.0, the lowest pH tested (Figure 1G). We next tested the ability of encapsulated DypB to degrade ABTS across a range of pH values. Similar to free DypB, the encapsulated enzyme had the highest activity at pH ~4.0 (Figure 1H). Taken together, these data demonstrate the stability and functionality of *M. tuberculosis* nanocompartments under proteolytic and acid stress, conditions that mimic the host lysosomal environment.

The Mtb DypB protein was previously shown to function as a bona fide dye-decolorizing peroxidase^9^. Because peroxidases consume H_2_O_2_, they often participate in defense against oxidative stress^16^. During macrophage infection, the phagocytes initiate an oxidative burst that exposes Mtb to H_2_O_2_^17^. We therefore reasoned that DypB-containing nanocompartments may function to protect Mtb from H_2_O_2_-induced stress. To test this hypothesis, we created a mutant strain lacking both genes from the nanocompartment operon (Figure 1A, Δoperon). Lysates from Δoperon mutants were used for nanocompartment purification (data not shown) and western blot analysis using an antibody for Cfp29 as a probe (Figure S1C). Data from these analyses revealed that Δoperon mutants cannot produce viable nanocompartments. In addition, we isolated a transposon mutant (*DypB::Tn*) containing an insertion in *DypB*, a mutation that eliminated expression of both Cfp29 and DypB (Figure S1D).

To test whether DypB-containing nanocompartments are required for defense against H_2_O_2_, we exposed wild-type and *DypB::Tn* Mtb to increasing concentrations of H_2_O_2_ and monitored bacterial survival. Mutants lacking DypB nanocompartments were more susceptible to H_2_O_2_ when compared with wild-type bacteria as measured by OD_600_ (Figure 2A) and by plating for CFU (Figure 2B). However, the phenotype was relatively modest. We next reasoned that DypB nanocompartments might be required to resist oxidative stress in acidic environments that more closely mimic the *in vivo* environment. During infection, Mtb bacilli encounter the low pH of the phagolysosome, and the ability to tolerate low pH is required for Mtb survival in both infected macrophages and mice^18^. We therefore tested the susceptibility of DypB mutants to a combination of H_2_O_2_ and acid stress (pH 4.5). Wild-type Mtb was able to withstand these conditions and did not significantly decrease in number over a three-day exposure period (Figure 2C). In contrast, the Δoperon mutant was resistant to each stressor individually, but was highly susceptible to H_2_O_2_ at pH 4.5 (Figure 2C). Thus, DypB-containing nanocompartments are required to protect bacteria from oxidative stress at low pH.

**Figure 2.**
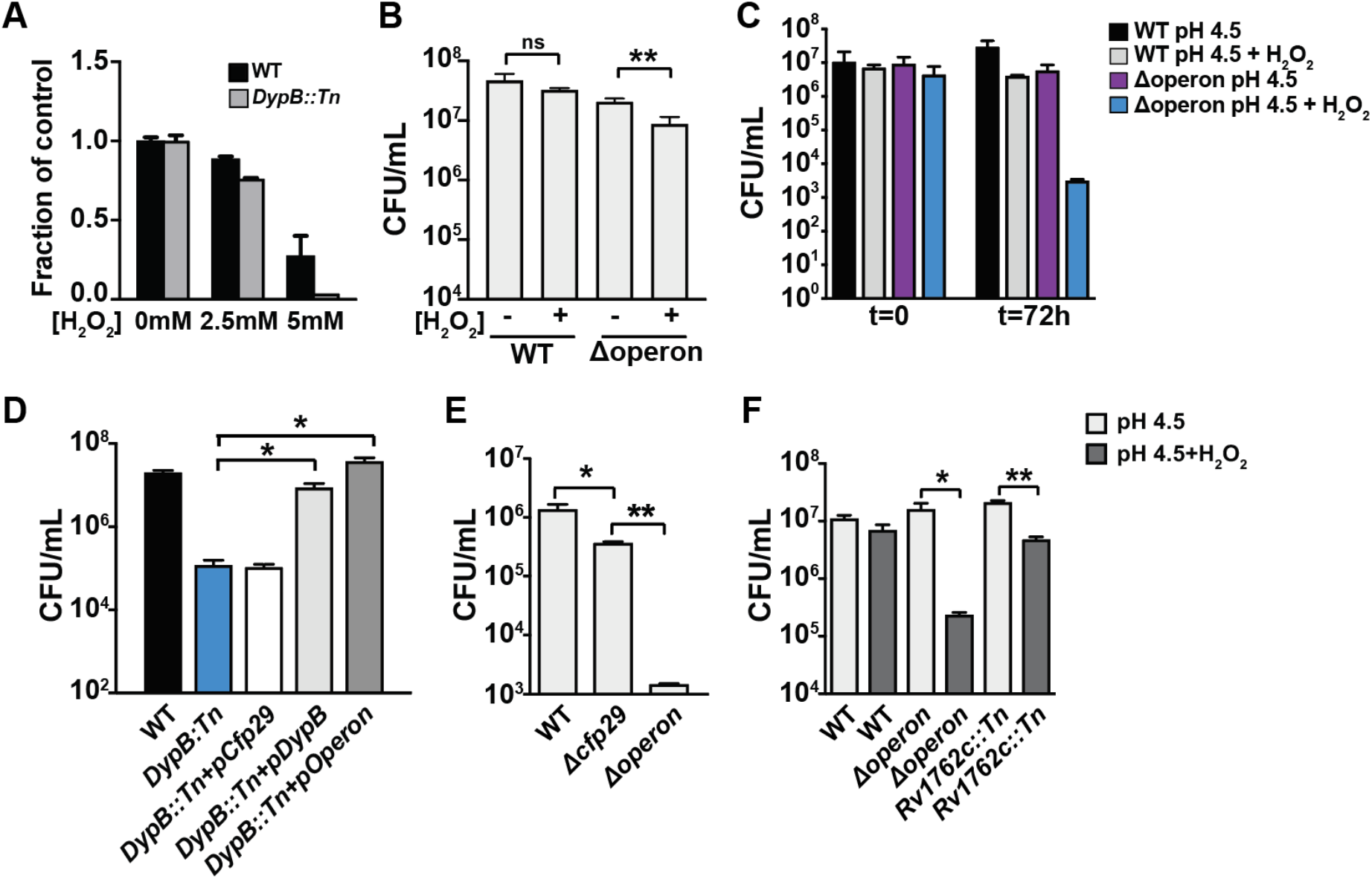
Nanocompartments protect Mtb from oxidative stress in acidic environments. **(A)** OD_600_ measurements of wild-type and *DypB::Tn* Mtb grown in 7H9 medium following exposure to H_2_O_2_ for 96 hours. Values reported are normalized to the untreated controls. CFU enumeration of wild-type and Mtb nanocompartment mutants grown in **(B)** standard 7H9 medium (pH 6.5) and **(C, E, F)** acidified 7H9 medium (pH 4.5) following exposure to oxidative stress (2.5 mM H_2_O_2_) for 72 hours. **(D)** CFU enumeration of wild-type, *DypB::Tn* mutants, and complemented mutants (pCfp29, pDypB, pOperon) following 24 hour exposure to oxidative stress (2.5 mM H_2_O_2_) in acidified 7H9 medium (pH 4.5). Figures are representative of at least 2 **(E,F)** or 3 **(A-D)** independent experiments. p-values were determined using unpaired t test. *p<0.05, **p<0.01.

We next sought to determine whether encapsulation of DypB is important for its activity in bacterial cells. To accomplish this, *DypB::Tn* mutants were complemented with constructs expressing DypB alone (pDypB), Cfp29 encapsulin alone (pCfp29), or both proteins (pOperon). As expected, expression of DypB did not restore production of nanocompartments, whereas expression of Cfp29 alone resulted in the formation of empty nanocompartment structures (Figure S1D). Co-expression of both proteins resulted in the formation of DypB-containing nanocompartments in the *DypB::Tn* mutant (Figure S1D). We exposed the full set of complemented mutants to H_2_O_2_ at pH 4.5 and determined survival after three days by plating for CFU. As expected, *DypB::Tn* was attenuated for survival under these conditions when compared to wild-type bacteria (Figure 2D). Restoring expression of the nanocompartment shell protein in the absence of DypB had no effect on bacterial survival (Figure 2D). Complementation by overexpression of unencapsulated DypB was sufficient to confer almost wild-type levels of resistance to oxidative and acid stress (Figure 2D). However, complementation with both Cfp29 and DypB resulted in enhanced resistance to these stressors when compared to mutants complemented with DypB alone (Figure 2B), suggesting that encapsulation of DypB enhances its function.

In the mutant complemented with DypB alone, it is possible that overexpression of the peroxidase compensates for the lack of encapsulation. We therefore examined the importance of DypB encapsulation in mutant strains in which we deleted *Cfp29* only (Δcfp29). Deletion of *Cfp29* resulted in a ~0.5 log decrease in bacterial viability after three days of exposure to H_2_O_2_ at pH 4.5, confirming that encapsulation of the peroxidase in the shell protein is important for full protection (Figure 2E). As expected, greater attenuation was observed in the mutant lacking both DypB and Cfp29 (Δoperon). Our mass spectrometry data suggested that an additional protein, Rv1762c, associates with the Mtb encapsulin nanocompartment (Figure 1E). To test whether Rv1762c has functional significance, we isolated a mutant with a transposon insertion in this gene (*Rv1762c::Tn*) and exposed it to H_2_O_2_ at pH 4.5. We found that *Rv1762c::Tn* is moderately susceptible to a combination of low pH and H_2_O_2_ (Figure 2F). Taken together, these data demonstrate that encapsulation of DypB enhances protection against oxidative stress in low pH conditions and that Rv1762c may have a functional role in the DypB encapsulin system.

The Mtb genome encodes ~18 putative peroxidase genes, including DypB. To determine whether other bacterial peroxidases participate in the defense against oxidative stress at low pH, we performed a transposon sequencing (Tn-seq) screen^19,20^. To do so, we created a transposon library containing ~100K individual transposon mutants and exposed this library to H_2_O_2_ at pH 4.5 for three days, at which point the surviving bacteria were plated. Transposon gene junctions were amplified and sequenced from the recovered bacteria and the sequencing data were analyzed using TRANSIT^21^ (Table S1). These data revealed that of the 18 putative enzymes encoded by Mtb that could be involved in oxidative defense, including peroxidases, catalases, and superoxide dismutases, only 2 genes were required for survival in the culture conditions—*DypB* and *KatG* (Figure 3A). *KatG* encodes a catalase-peroxidase that is important for defense against oxidative stress in the host^22,23^. Coincidentally, KatG is also required for activation of isoniazid (INH), a central component of anti-TB therapy. By contrast, DypB did not detectably react with INH (data not shown). Approximately 10% of TB cases are caused by INH-resistant bacteria, many of which have loss-of-function mutations in KatG^24^. Importantly, KatG variants that do not activate INH often display a concomitant decrease in peroxidase and catalase activity^25–27^. Both the high proportion of KatG mutant bacteria in the human population and studies of virulence in animals suggest that KatG mutants are still capable of growth *in vivo*^23,28^. The fact that both DypB and the KatG mutants are important for survival in oxidative stress at low pH suggests that the DypB nanocompartment may provide redundancy that compensates for a loss of KatG in INH resistant strains of Mtb.

**Figure 3.**
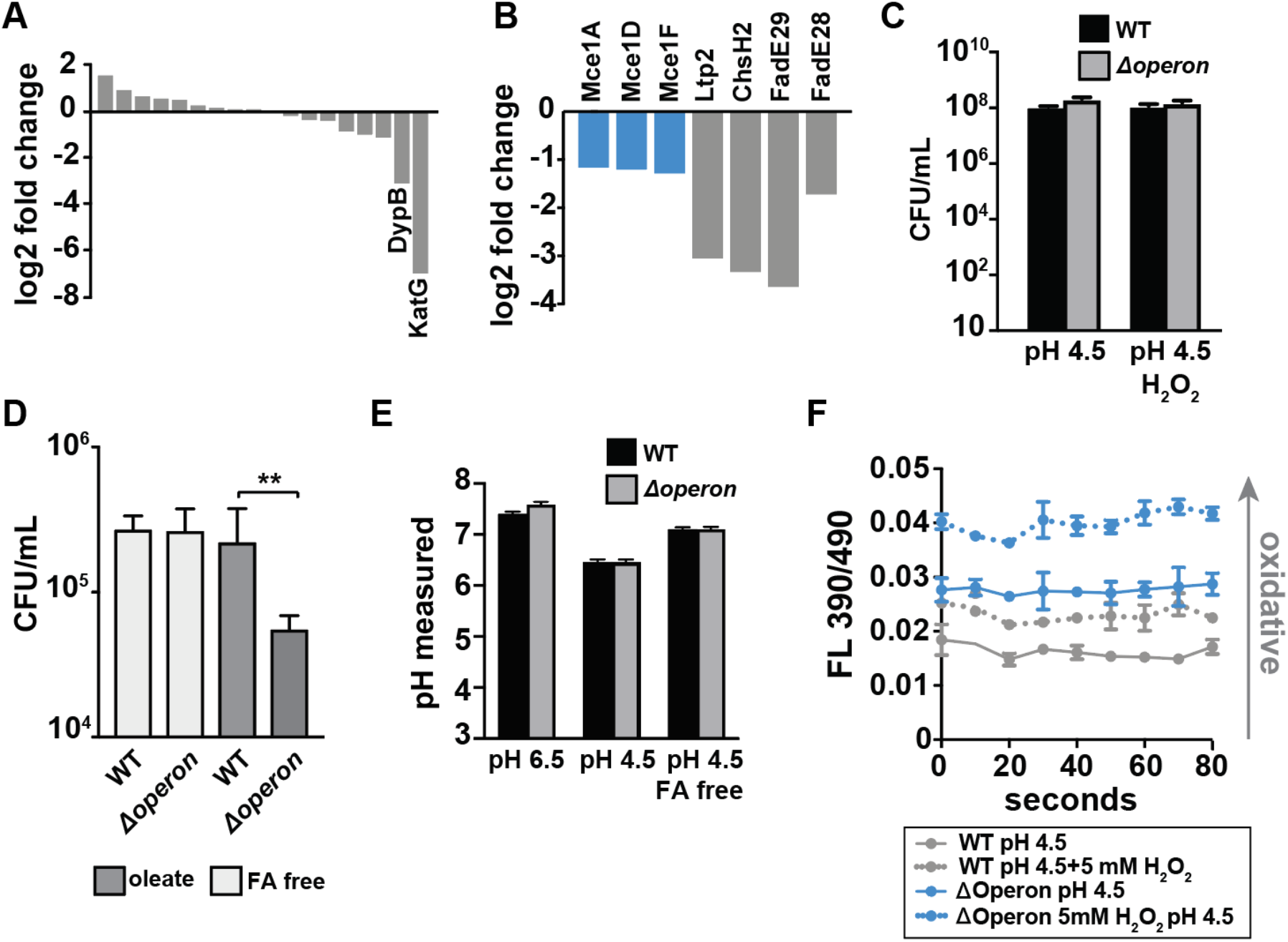
Susceptibility of Mtb nanocompartment mutants to oxidative and acid stress is mediated by free fatty acids. **(A)** Tn-seq data showing normalized sequence reads per gene for all putative Mtb peroxidases and **(B)** lipid and cholesterol metabolism Mtb mutants that were significantly attenuated following 72 hour exposure to 2.5 mM H_2_O_2_ at pH 4.5. **(C)** CFU enumeration of wild-type Mtb and Δoperon mutants following 24 hour exposure to 2.5 mM H_2_O_2_ at pH 4.5 in Sauton’s minimal medium and **(D)** 72 hour exposure to 2.5 mM H_2_O_2_ at pH 4.5 in 7H9 medium prepared using fatty-acid (FA) free BSA +/− oleic acid (150 uM). **(E)** Intrabacterial pH measurements of wild-type and Δoperon Mtb expressing pUV15-pHGFP following 20 minute exposure to 5 mM H_2_O_2_ at pH 6.5 or pH 4.5. 7H9 medium was prepared with standard BSA or FA free BSA. **(F)** Fluorescence emissions of wild-type and Δoperon Mtb expressing mrx1-roGFP exposed to 5 mM H_2_O_2_ at pH 4.5 in 7H9 medium for 20 minutes. Data are reported as a ratio of fluorescence emissions following excitation at 490 nm and 390 nm. Figures are representative of at least 2 **(D)** or 3 **(A-C; E-F)** independent experiments. p-values were determined using an unpaired t test. *p<0.05, **p<0.01.

In the Tn-seq data, we observed that mutants involved in lipid or cholesterol metabolism were also attenuated when exposed to H_2_O_2_ at pH 4.5 (Figure 3B). *M. tuberculosis* biology is highly linked to lipid biology; Mtb granulomas are rich in free fatty acids^29^ and the bacteria utilize lipids as a source of nutrition both during macrophage infection *ex vivo* and growth *in vivo*^30–32^. Based on the Tn-seq data, we hypothesized that lipids may mediate the sensitivity of DypB mutants to oxidative stress under acidic conditions. The standard Mtb culture medium, 7H9, contains Bovine Serum Albumin (BSA), which has binding sites for fatty acids and may also serve as a cholesterol shuttle in serum^33,34^. We tested whether Δoperon mutants were susceptible to acid and oxidative stress in Sauton’s broth, a minimal medium that lacks BSA, and found that Δoperon mutants persisted to the same degree as wild-type bacteria (Figure 3C). To test whether lipids bound to BSA mediate the susceptibility of DypB mutants to acid and oxidative stress, we cultured wild-type and Δoperon Mtb in media constituted with fatty acid-free BSA and exposed the bacteria to H_2_O_2_ at pH 4.5 for three days. In the absence of BSA bound lipids, we found that the Δoperon mutant was not susceptible to oxidative stress at low pH (Figure 3D). 7H9 medium prepared with lipid free BSA was then reconstituted with oleic acid, an abundant lipid found in mammalian systems^35^. Reconstitution of fatty acid free medium with oleic acid restored toxicity to nanocompartment mutants under conditions of acid and oxidative stress (Figure 3D). Taken together, these data suggest that the phagosomal lipids used by Mtb as a carbon source may be toxic to bacteria lacking functional nanocompartments under conditions of oxidative stress

We considered that lipids might disrupt Mtb membranes, resulting in altered pH homeostasis, and a drop in cytosolic pH. Interestingly, in acidified medium we found that the intracellular pH of wild-type bacteria dropped from ~pH 7.5 to ~ pH 6.2 and this decrease was dependent on the presence of albumin-bound fatty acids (Figure 3E). These data confirm published results demonstrating that the cytosolic pH of Mtb drops significantly during acid exposure.^18^ A lower cytosolic pH would possibly impair the function of many other bacterial enzymes important for resisting oxidative stress and place increasing importance on enzymes that can function in acidic environments, such as DypB.^23^ Taken together, these data may explain why functional nanocompartments are critical for bacterial survival when Mtb is exposed to acid and oxidative stress in a fatty acid-rich environment.

We next sought to test whether nanocompartment mutants have altered redox homeostasis in the presence of oxidative stress. We transformed wild-type and mutant strains with Mrx1-roGFP, a fluorescent reporter of redox potential in mycobacteria^36^. Remarkably, the Δoperon mutant had an intracellular environment at baseline that was more oxidizing than wild-type bacteria exposed to H_2_O_2_ (Figure 3F), indicating a failure of redox homeostasis in mutants lacking the DypB encapsulin system. This oxidizing cellular environment was further enhanced by treatment with H_2_O_2_ (Figure 3F). Thus, mutants lacking DypB-containing nanocompartments exhibit significant dysregulation of redox homeostasis.

Our *in vitro* findings that DypB nanocompartments are important for defense against a combination of oxidative stress, low pH, and fatty acids suggested that this system may be important for Mtb survival in the phagosomal environment. Therefore, we sought to determine whether DypB nanocompartments are required for bacterial growth in host cells. We found that both the *DypB::Tn* and Δoperon mutants had impaired growth in murine bone-marrow derived macrophages that was manifest by two days after infection (Figure 4A, 4B). One of the antibiotics used to treat TB infection, Pyrazinamide (PZA), is most effective in low pH environments, such as the phagolysosome. PZA is thought to disrupt the cell wall of Mtb, which leads to acidification of the bacterial cytosol. A recent study in human TB patients demonstrated that the pH of necrotic lung cavities is ~5.5, a finding that may also explain the efficacy of pyrazinamide in treating Mtb infection^37^. Since DypB is critical for protection of Mtb against oxidative stress at low pH, we considered whether nanocompartments may mediate Mtb resistance to PZA. To test bacterial susceptibility to PZA, we treated wild-type and Δoperon Mtb with PZA in the presence of H_2_O_2_ at pH 5.5. We found that mutants lacking the DypB nanocompartment were sensitive to concentrations of PZA that did not impact growth of the wild-type bacteria (Figure 4C).

**Figure 4.**
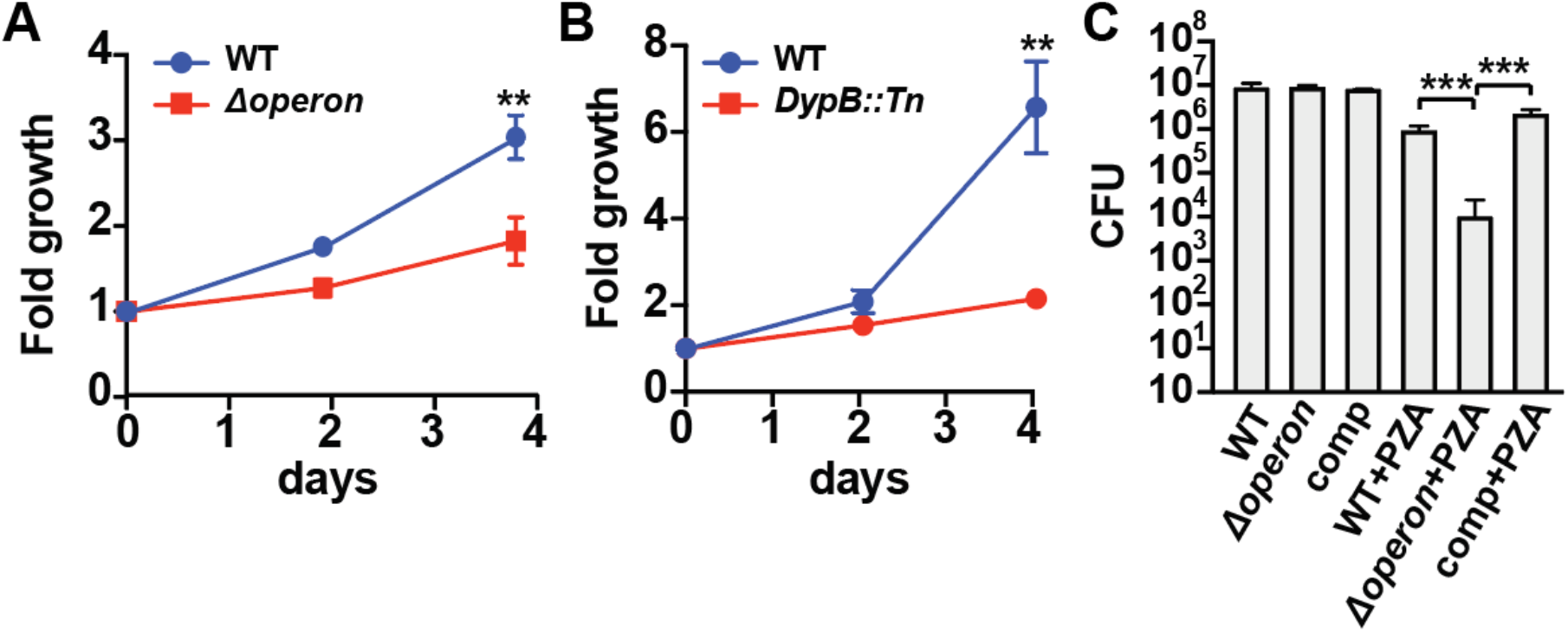
Nanocompartment mutants are attenuated for survival in macrophages and are more susceptible to pyrazinamide treatment. CFU enumeration of wild-type Mtb and **(A)** Δoperon or **(B)** DypB::Tn mutants during infection of murine bone marrow-derived macrophages. Macrophages were infected with a bacterial MOI of 1 and CFUs were enumerated immediately following phagocytosis and at days 2 and 4. Error bars are SD from 4 replicate wells. **(C)** CFU enumeration of wild-type and Mtb Δoperon mutants following 72 hour exposure to pyrazinamide (24 μg/mL) and H_2_O_2_ (2.5 mM) in acidified 7H9 medium (pH 5.5). Comp=*Δoperon+pOperon*. Figures are representative of at least 2 independent experiments; p-values were determined using an unpaired t test. **p<0.01; ***p<0.001.

Encapsulin nanocompartments are a largely uncharacterized proteinaceous organelle widespread in bacteria and archaea. Here we provide some of the first functional data demonstrating the physiological significance of a nanocompartment and its cargo in any organism. Furthermore, our finding that encapsulation of DypB promotes its function in resisting oxidative stress represents the first demonstration that encapsulation can promote the function of an encapsulated enzyme in an endogenous setting. Furthermore, we have demonstrated that an encapsulin system is important for resisting oxidative stress in the context of a globally significant pathogen, *Mycobacterium tuberculosis,* expanding our understanding of how this pathogen combats the hostile *in vivo* environment.

## Materials and Methods

### Ethics statement

All procedures involving the use of mice were approved by the University of California, Berkeley Institutional Animal Care and Use Committee (protocol no. R353-1113B). All protocols conform to federal regulations, the National Research Council’s *Guide for the Care and Use of Laboratory Animals*, and the Public Health Service’s *Policy on Humane Care and Use of Laboratory Animals*.

### Mycobacterium tuberculosis bacterial strains and plasmids

The *M. tuberculosis* strain H37Rv was used for all experiments. The transposon mutants DypB::Tn and Rv1762c::Tn were picked from an arrayed transposon mutant library generated at the Broad Institute. The ΔOperon, ΔCfp29, and ΔDypB strains were made by homologous recombination using the pMSG361 vector^38^. For genetic complementation studies, the region encoding GFP and KanR in pUV15tetORm^39^ was substituted via GoldenGate cloning with open reading frames for Rv0798c, Rv0799c, or the whole nanocompartment operon (Rv0798-99c). Expression of the complementation constructs was induced with anhydrotetracycline (200 ng/mL). To measure redox homeostasis, strains were transformed with pMV762-mrx1-roGFP2^36^. To measure intrabacterial pH, strains were transformed with pUV15-pHGFP (Addgene). The transposon mutant library for Tn-Seq was generated in *M. tuberculosis* using the ΦMycoMarT7 transposon donor plasmid.

### *M. tuberculosis* bacterial cell culture

For infections*, M. tuberculosis* was grown to mid-log phase (OD_600_ = 0.5-1.0) in Middlebrook 7H9 liquid medium supplemented with 10% albumin-dextrose-saline, 0.4% glycerol, and 0.05% Tween-80 or on solid 7H10 agar plates supplemented with Middlebrook OADC (BD Biosciences) and 0.4% glycerol. When specified, Tween-80 was substituted with 0.05% Tyloxapol, and 10% albumin-dextrose-saline was prepared with fatty acid free BSA (Sigma-Aldrich). Sauton’s media was prepared with tyloxapol as previously specified^40^.

### DypB activity assays

Activity of the encapsulated and unencapsulated DypB was performed using methods adapted from Contreras *et al.* 2014. Briefly, DypB concentration for the encapsulated and unencapsulated DypB was determined by absorbance of the heme prosthetic group at 411 nm. Reactions were performed using 5 nM DypB, 480 nM H_2_O_2_, and 480 nM 2,2’-azino-bis (3-ethylbenzothiazoline-6-sulfonic acid) (ABTS) in 100mM sodium citrate buffer pH 4-6. Product formation was monitored over 20 minutes via absorbance at 420 nm using a Varian Cary® 50 UV-Vis Spectrophotometer (Agilent).

### Nanocompartment Purification from Mtb

For each purification, 1.5 L of *M. tuberculosis* was grown to mid-log phase in standard 7H9 and washed with PBS. Bacteria were pelleted and lysed in buffer by bead beating (for 50 mL of buffer, PBS with 1 mM PMSF was supplemented with 50 mg lysozyme, 20 U DNaseI, and 100 μg RNase A). Lysates were passaged twice through 0.2 μm filters before removal from the BSL3. Clarified lysates were prepared by centrifugation at 20,000 x *g* for 20 minutes in a JA-20 rotor. Following clarification, lysates were layered onto top of a 38% sucrose cushion and centrifuged for 18 hours at 100,000 x *g* in a type 50.2 Ti rotor. The supernatant was discarded and the pellet was resuspended in 200 *μ*L of PBS. Resuspended pellets were layered on top of a 10-50% sucrose gradient and centrifuged for 21 hours at 100,000 x g in a SW 41 Ti rotor. The gradient was fractionated and aliquots from each fraction were analyzed by SDS-PAGE for the presence of Cfp29.

### Expression of Holo-nanocompartment and naked DypB in *E. coli*

Plasmids for the expression of the Holo-nanocompartment (DypB-loaded) and naked DypB constructs were designing using Gibson Assembly (NEB). Each construct was cloned into a pET-14-based destination vector containing a T7 promoter. The naked DypB construct contained an N-terminal poly-histidine tag for affinity purification. These constructs were transformed into *E. coli* BL21 (DE3) LOBSTR cells for protein overexpression. Cells were grown in LB media containing 60 μg/mL kanamycin at 37°C with shaking at 250rpm until cultures reached an optical density (OD_600_ = 0.5-0.6). Samples were then induced with 0.5 mM IPTG and grown overnight at 18°C. Liquid cultures were harvested by centrifugation at 5000 x *g* for 20 minutes at 4°C, flash frozen in liquid nitrogen, and then stored at −80°C for future use.

### Purification of Holo-nanocompartment complex from *E. coli*

Cell pellets (5 g dry cell mass) were thawed at room temperature and resuspended in 50 mL of lysis buffer (20 mM Tris-HCl pH 8, 150 mM NH_4_Cl, 20 mM MgCl_2_) supplemented with 50 mg lysozyme, 20 U DNaseI, 100 ug RNase A. Samples were lysed by three passages through an Avestin EmulsiFlex-C3 homogenizer and clarified via centrifugation (15,000 x g, 30 min, 4°C). The clarified lysate was then spun at 110,000 x *g* for 3 hours at 4 °C. The supernatant was discarded and the resulting pellet was resuspended with wash buffer (20 mM Tris pH 8, 150 mM NH_4_Cl, 20 mM MgCl_2_ supplemented with 1X Cell Lytic B (Sigma-Aldrich). The sample was then spun at 4,000 x *g* at 4°C for 10 min followed by removing the supernatant and resuspension of the pellet in 4 mL of 50 mM Tris-HCl pH 8, 300 mM NaCl. The sample was then incubated at room temperature for 10 minutes to allow for solubilization and then centrifuged at 4000 x *g* at 4°C for 10 minutes to remove insoluble material. The resulting supernatant was then concentrated using Vivaspin® 6 100,000 MWCO concentrator columns (Sartorius). The sample was then purified via size exclusion chromatography using a Superose™ 6 Increase column (GE Life Sciences) and fractions were analyzed by SDS-PAGE using 4-20% Criterion™ polyacrylamide gels (Bio-Rad) and visualized with GelCode Blue stain (ThermoFisher).

### Purification of unencapsulated DypB from *E. coli*

Cell pellets (5 g dry cell mass) were thawed at room temperature and resuspended in 50 mL of buffer A (25 mM Tris HCl pH 7.5, 150 mM NaCl, 20 mM imidazole) supplemented with 50 mg lysozyme, 20 U DNaseI, 100 μg RNase A. Samples were lysed by three passages through an Avestin EmulsiFlex-C3 homogenizer and clarified via centrifugation (15,000 x g, 30 min, 4°C). The resulting supernatant was then bound to HisPur™ Ni-NTA resin (ThermoFisher Scientific) for 90 minutes at 4°C and then applied to a gravity column. The nickel resin was then washed with 30 resin volumes of buffer B (25 mM Tris-HCl pH 7.5, 150 mM NaCl, 40mM imidazole) prior to eluting with buffer C (25 mM Tris-HCl pH 7.5, 150 mM NaCl, 350 mM imidazole). The eluate was then concentrated using Vivaspin® 20 10,000 MWCO concentrator columns (Sartorius) and desalted into 25mM Tris pH 8, 300mM NaCl using Econo-Pac®10DG desalting columns (BioRad). The SUMO tag was removed upon addition of SUMO protease at a 1: 300 (SUMO protease: DypB) molar ratio and incubating overnight at 4°C. Purification was finished by size exclusion chromatography with a Superose™ 6 Increase column (GE Life Sciences).

### Negative stain Transmission Electron Microscopy

Nanocompartment samples were diluted to 50 nM and applied to Formvar/ carbon-coated copper grids. The grids were then washed with MilliQ water three times followed by staining with 2% (w/v) uranyl acetate. Grids were examined using the FEI Tecnai 12, 120kV transmission electron microscope, and images were captured with a charge-coupled device (CCD) camera.

### Cathepsin B proteolysis assay

Proteolysis was performed using 2 μM of encapsulated or unencapsulated DypB in proteolysis buffer (100mM Sodium acetate pH 5, 2mM DTT) with or without the addition of 1 μM Cathepsin B (Sigma-Aldrich Cat# 219362). Reactions were run for 14 hours at 40°C followed by analysis of samples by SDS-PAGE using 4-20% Criterion™ polyacrylamide gels (Bio-Rad) and visualized with GelCode Blue stain (ThermoFisher).

### Exposure to oxidative and pH stress

*M. tuberculosis* was grown to mid-log phase in 7H9 media. Bacteria were diluted to OD_600_ = 0.1 in 10 mL of specified media at pH 4.5-6.5 and H_2_O_2_ was added to bacterial cultures at specified concentrations. Bacteria were incubated with stressors for 24 or 72 hours. CFUs were enumerated by diluting bacteria in PBS with 0.05% Tween-80 and plating serial dilutions on 7H10 agar.

### Measurement of redox homeostasis

*M. tuberculosis* strains were transformed with a plasmid expressing mrx1-roGFP2 and grown to mid-log phase in 7H9. Bacteria were diluted to OD_600_ = 0.25 in 200 μL of specified media and added to 96 well plates. Upon addition of H_2_O_2_ (5 mM), fluorescent emissions were recorded at 510 nm after excitation at 390 nm and 490 nm using a Spectramax M3 spectrophotometer. Values reported are emissions ratios (390 nm/ 490 nm) and were measured 20 minutes following addition of H_2_O_2_.

### Measurement of intrabacterial pH

*M. tuberculosis* strains were transformed with a plasmid expressing pHGFP and grown to mid-log phase in 7H9. To prepare standards, 1.5 × 10^8 bacterial cells were pelleted and resuspended in 400 μL lysis buffer (50 mM Tris-HCl pH 7.5, 5 mM EDTA, 0.6% SDS, 1 mM PMSF) before bead beating. Cell debris were pelleted and clarified lysates were kept at 4°C until use, at which point 10 μL of clarified lysate were added to 200 μL of medium with varying pH levels (4.5-8.0). To prepare samples, 1.5 × 10^8 bacterial cells were pelleted and washed with PBS twice before being resuspended in specified media and diluted to OD_600_ = 0.5 in 200 μL of medium and added to 96 wells plates. Upon addition of H_2_O_2_ (5 mM), fluorescent emissions were recorded at 510 nm following excitation at 395 nm and 475 nm. Values reported were interpolated from 395/475 ratios obtained from the standard curve.

### Western blot analysis of Cfp29 expression

*M. tuberculosis* strains were grown to mid-log phase in 7H9 medium. Bacteria were pelleted and washed twice with PBS prior to resuspension in lysis buffer (50 mM Tris-HCl pH 7.5, 5 mM EDTA, 0.6% SDS, 1 mM PMSF). Samples were lysed using a bead-beater, and cell debris were pelleted. Clarified lysates were heat-sterilized at 100°C for 15 minutes and frozen prior to use. Total protein lysates were analyzed by SDS-PAGE using precast Tris-HCl 4-20% criterion gels (Bio-Rad). Primary polyclonal antibodies for Cfp29 were generated by GenScript USA Inc via immunization of rabbits with three peptides from the protein sequence. HRP-conjugated goat anti-rabbit IgG secondary antibodies were used (sc-2030; Santa Cruz Biotechnology). Western Lightning Plus-ECL chemiluminescence substrate (Perkin Elmer) was used and blots were developed using a ChemiDoc MP System (Bio-Rad).

### Infection of murine macrophages

Macrophages were derived from bone marrow of C57BL/6 mice by flushing cells from femurs. Cells were cultured in DMEM supplemented with 10% FBS and 10% supernatant from 3T3-M-CSF cells for 6 days, with feeding on day 3. After differentiation, BMDMs continued to be cultured in BMDM media containing M-CSF. For infection, BMDMs were seeded at a density of 5 × 10^4 cells per well in a 96-well dish. BMDMs were allowed to adhere overnight and then infected with DMEM supplemented with 5% FBS and 5% horse serum (BMDMs) at a multiplicity of infection of 1. Following a 4-hour phagocytosis period, infection medium was removed and cells were washed with room temperature PBS before fresh, complete medium was added. For CFU enumeration, medium was removed and cells were lysed in water with 0.5% Triton-X and incubated at 37°C for 10 minutes. Following the incubation, lysed cells were resuspended and serially diluted in PBS with 0.05% Tween-80. Dilutions were plated on 7H10 plates

### Transposon-sequencing screen

A transposon mutant library in H37Rv was grown to mid-log phase in 7H9. Bacteria were diluted to OD_600_ = 0.1 in 10 mL 7H9 at pH 4.5 with 2.5 mM H_2_O_2_. Mutants were exposed to these stressors for 72 hours and then diluted to 15,000 CFU/mL in PBS with 0.05% Tween-80. Approximately 30 thousand bacteria were plated onto six 245 mm × 245 mm 7H10 plates supplemented with 0.05% Tween-80 and Kanamycin (50 μg/mL). Control libraries were not exposed to low pH or H_2_O_2_ and were plated onto 7H10 plates. Colonies grew for 21 days and were collected for genomic DNA isolation. Samples for sequencing were prepared by the University of California, Davis Genome Center DNA Technologies Core by following the protocol outlined by Long et al^41^. PE100 reads were run on an Illumina HiSeq with ~20 million reads per sample. Sample alignment and TRANSIT pre-processing were performed by the University of California, Davis Bioinformatics Group as previously outlined^21^. TRANSIT analysis was performed as specified by DeJesus et al^21^. Resampling analysis was preformed using the reference genome H37RvBD_prot and the following parameters: for global options, 0% of the N- and C-terminus were ignored; for resampling options, 10,000 samples were taken and normalized using the TTR function. Correction for genome positional bias was performed. Statistical significance was determined by p value ≤ 0.05 and log2 fold change ≤-1 or by p-adjusted value ≤ 0.05.

## Acknowledgments

We thank Tom Ioerger and Michael DeJesus for assistance with TRANSIT analysis and Amit Singh for the kind gift of pMV762-mrx1-roGFP2. We thank Jeff Cox and members of the Cox lab for helpful discussions. This work was supported by funding from the Center for Emerging and Neglected disease for funding to KAL, the National Institute of General Medical Sciences (R01GM129241) to DFS and the National Institute of Allergy and Infectious Diseases (1R01AI143722) to SAS.

---

**Figure S1.**
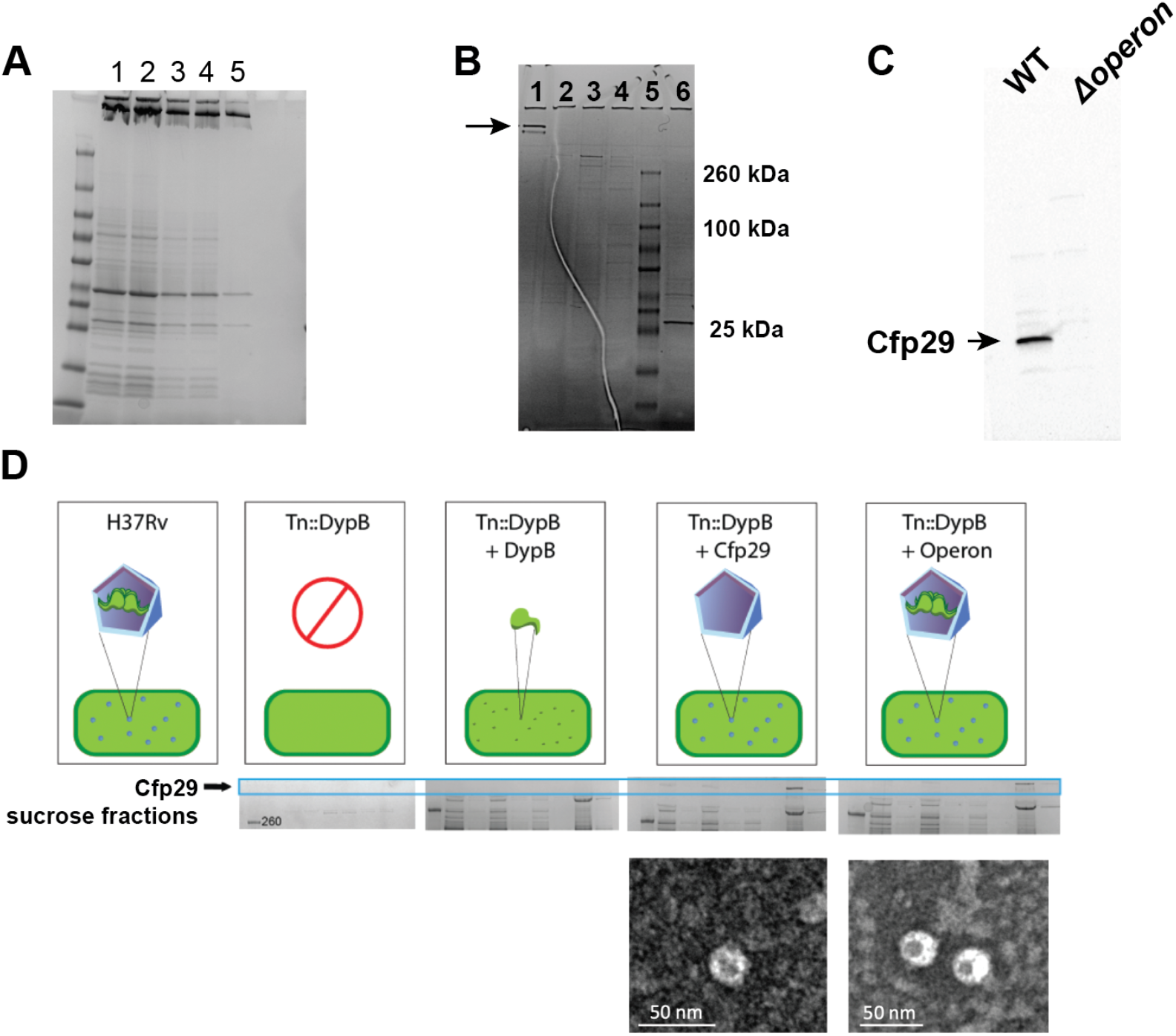
**(A)** Coomassie stained SDS-PAGE of fractions collected during purification of nanocompartments heterologously expressed in *E. coli*: (1) Ultracentrifugation pellet post CellLytic B solubilization (2) Size exclusion chromatography input (3) lane 1 diluted (4) lane 2 diluted (5) encapsulin fraction from size exclusion. **(B)** Coomassie stained SDS-PAGE of sucrose fractions collected during purification of nanocompartments from wild-type Mtb lysates: (1) Fraction containing assembled encapsulin nanocompartment complexes (5) Ladder (6) Lane 1 boiled in SDS for 30 minutes to dissociate encapsulin nanocompartment into monomers. **(C)** Western blot for Cfp29 from wild-type Mtb (lane 1) and Δoperon mutant (lane 2) lysates. **(D)** Complementation strategy schematic for *DypB::Tn* mutants (top). *DypB::Tn* mutants were transformed with ATc-inducible complementation constructs encoding the unencapsulated cargo protein (pDypB), the encapsulin shell protein (pCfp29), or the nanocompartment operon (pOperon). Lysates from each strain were used for nanocompartment purification. Sucrose fractions containing high molecular weight Cfp29 protomers were identified in complemented strains expressing the encapsulin shell and the operon (middle) and were analyzed using TEM (bottom).

**Figure S2.**
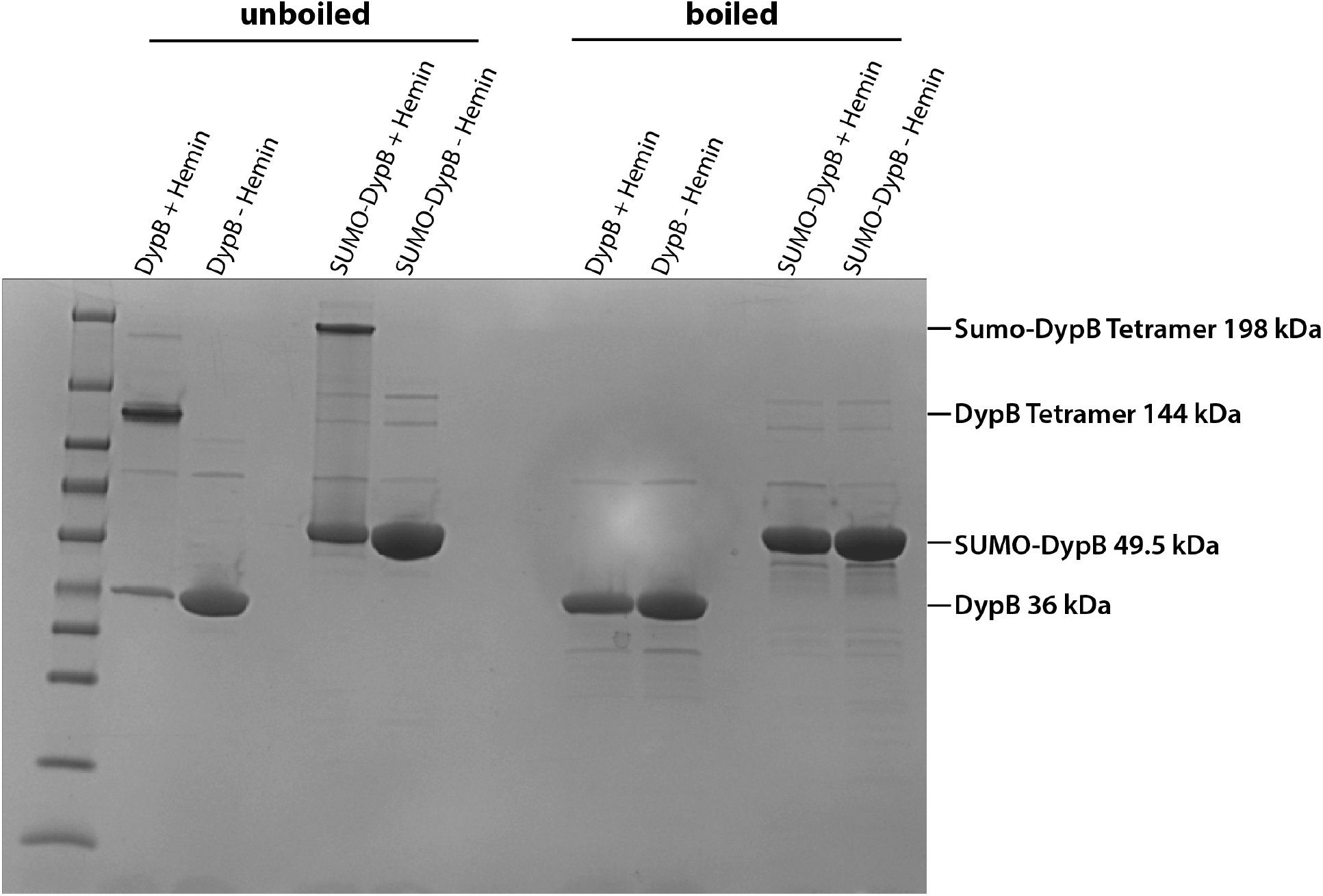
SDS-PAGE analysis of DypB purified from *E. coli*. DypB samples with or without addition of hemin were analyzed by SDS-PAGE. Samples were loaded either in their unboiled native state (left half) or heat-denatured by boiling at 95°C for 15 minutes. Addition of hemin yields a tetrameric DypB at 144 kDa.

